# Predation by a ciliate community mediates temperature and nutrient effects on a peatland prey prokaryotic community

**DOI:** 10.1101/2024.04.05.588366

**Authors:** Katrina DeWitt, Alyssa A. Carrell, Jennifer D. Rocca, Samantha Votzke, Andrea Yammine, Ariane L. Peralta, David J. Weston, Dale A. Pelletier, Jean P. Gibert

**Author notes:** Address correspondence to Katrina DeWitt. Samantha Votzke, Department of Earth and Planetary Sciences, Johns Hopkins University, Baltimore, MD, USA.

## Abstract

Temperature significantly impacts microbial communities’ composition and function, which plays a vital role in the global carbon cycle that determines climate change. Nutrient influxes often accompany rising temperatures due to human activity. While ecological interactions between different microorganisms could shape their response to environmental change, we do not understand how predation may influence these responses in a warmer and increasingly nutrient-rich world. Here, we assess whether predation by a ciliate community of bacterial consumers influences changes in the diversity, biomass, and function of a freshwater prokaryotic community under different temperature and nutrient conditions. We found that predator presence mediates the effects of temperature and nutrients on total prokaryotic community biomass and composition through various mechanisms, including direct and indirect effects. However, the total community function was resilient. Our study supports previous findings that temperature and nutrients are essential drivers of microbial community composition and function but also demonstrates how predation can mediate these effects, indicating that the biotic context is as important as the abiotic context to understanding microbial responses to novel climates.

**Importance:** While the importance of the abiotic environment in microbial communities has long been acknowledged, how prevalent ecological interactions like predation may influence these microbial community responses to shifting abiotic conditions is largely unknown. Our study addresses the complex interplay between temperature, nutrients, predation, and their joint effects on microbial community diversity and function. Our findings suggest that while temperature and nutrients are fundamental drivers of microbial community dynamics, the presence of predators significantly alters these responses. Our study underscores the impact of abiotic factors on microbial communities and the importance of accounting for the biotic context in which these occur to understand, let alone predict, these responses properly.

## Introduction

Rising global temperatures are currently affecting populations^1,2^, communities^3^, and ecosystems^4^ rising organismal metabolic rates^5,6^, leading to higher energetic demands from populations^7–9^, and cascading effects within communities and ecosystems^5,10–12^. Additionally, human activity is increasing nutrient loads and mineralization rates worldwide^13^, which also affects communities and ecosystems^14–19^. While the independent effects of temperature and nutrients are relatively well-understood^20–22^, their joint increase can interactively influence communities^23–28^, and these interactions can be hard to both understand and predict. Communities of prokaryotes, in particular, comprise upwards of 14% of all existing biomass on Earth^29^ are present on all continents and ecosystems^30,31^, and play a central role in the global carbon and nutrient cycles^32–35^. As with other organisms, prokaryotic respiration often increases with temperature^5–7,36–39^, possibly leading to a "warming begets warming" scenario. Increasing nutrient loads can also increase heterotrophy and respiration^14,15^ in these communities. However, how rising temperature and nutrients might jointly influence prokaryotes in an increasingly warm, nutrient-rich, and human-dominated world is unclear.

Prokaryotic communities might also respond to the rapid rewiring^40–42^ of the broader food webs they are a part of^43^. Indeed, bacterivores are predicted to increase foraging rates to offset increased metabolic costs^44^, possibly resulting in decreased prokaryotic biomass with warming^45–48^, which could, in turn, impact the composition and function of prokaryotic communities in future climates^48–52^. Ciliate microbes are among the most abundant predators of prokaryotes worldwide^29,30,53,54^. These microbial predators show strong temperature^39,55–57^ and nutrient responses^28,58–60^, temperature-dependent population dynamics^39,57^ and feeding interactions^6,56^, and predation by a single ciliate has been shown to influence prokaryotic community structure, dynamics, and function^61,56,62,28^. Prokaryotic communities will interact with ciliate communities in nature –not just a single species– but how the presence of a ciliate community might influence the response to joint changes in temperature and nutrients is not known.

Here, we examine whether and how the presence of a ciliate community on a mostly prokaryotic prey community influences how nutrient additions and temperature affect total microbial biomass, diversity, composition, and function. To do so, we used experimental microcosms containing a synthetic community of naturally co-occurring ciliates^63^ and a peatland prokaryotic community to address: 1) How do temperature, nutrients, and the presence of a predatory ciliate community jointly affect prokaryotic community biomass, diversity, and composition? 2) What are the consequences for total microbial community respiration (i.e., function)? However, changes in the prey prokaryotic community might give feedback to the ciliates, and the ciliates might also directly respond to temperature and nutrients. So, we also ask: 3) How do temperature and nutrients affect the ciliates? We combined these answers to provide a perspective on how prokaryotes respond directly to temperature and nutrients and indirectly through ciliate responses to those same factors and their resulting impacts on the prokaryotes.

We hypothesize that prokaryotic biomass and total community respiration rates should increase with temperatures and nutrients^64,65^. Predation by a single ciliate decreases prokaryotic biomass^66,67^, so predation by the ciliate community should also lead to lower prokaryotic biomass, resulting in lower total respiration rates (prokaryotes + ciliates)^45^. However, ciliate presence can sometimes facilitate prokaryotic growth by mobilizing otherwise inaccessible resources (e.g., fertilization)^68,69^. Alternatively, the presence of a ciliate community could increase prokaryotic biomass and total community respiration rates. Ciliate presence should result in a compositional shift among the prokaryotes due to the differential consumption or facilitation^66,67^. Last, predation by a single ciliate species can interact with temperature and nutrients, to co-determine total respiration and biomass responses of the prokaryotes^52^, and so we expect that an entire community of ciliates might have similar effects. Because, the ciliates can themselves show ecological^28^ and phenotypic^70,57^ responses to temperature and nutrients^71,72^, we hypothesize that the prokaryotic community likely will respond to both direct, and indirect effects of temperature and nutrients, the later mediated by the ciliate direct responses to temperature and nutrient change.

## Results

### Direct effects of temperature, nutrients, and ciliate community on prokaryotic biomass and total community respiration

We conducted a fully factorial microcosm experiment manipulating temperature (22°C/25°C), nutrient availability (low/high), and the presence of an 8-species synthetic ciliate community known to co-occur in nature. After three weeks, we quantified prokaryotic biomass via spectrophotometry (OD600) and total community respiration using real-time respirometry (see Methods).

Contrary to what was observed in mono-ciliate treatments^52^, the same approach we used, we observed an increase in prokaryotic biomass in the presence of the ciliate community (effect = 0.151 ± 0.056 SE, *t* = 2.677, *df* = 111, *p* = 0.009; Figure 1A). This effect was temperature-dependent, as indicated by a significant negative interaction between temperature and ciliate presence (interaction = –0.0067 ± 0.0024 SE, *t* = –2.783, *df* = 111, *p* = 0.006; Figure 1B, 1C). Additionally, ciliates and nutrients interacted: under low nutrient conditions, the presence of ciliates reduced prokaryotic biomass (interaction = –0.206 ± 0.080 SE, *t* = –2.577, *df* = 111, *p* = 0.011; Figure 1D, 1E). Finally, we detected a significant three-way interaction between temperature, nutrient level, and ciliate presence (interaction = 0.0092 ± 0.0034 SE, *t* = 2.700, *df* = 111, *p* = 0.008), suggesting that the temperature effect on biomass depends on both nutrient availability and protist presence (Figure 1B, 1C). In contrast, the main effects of temperature (*p* = 0.986) and nutrient treatment (*p* = 0.787) were not significant.

**Figure 1.**
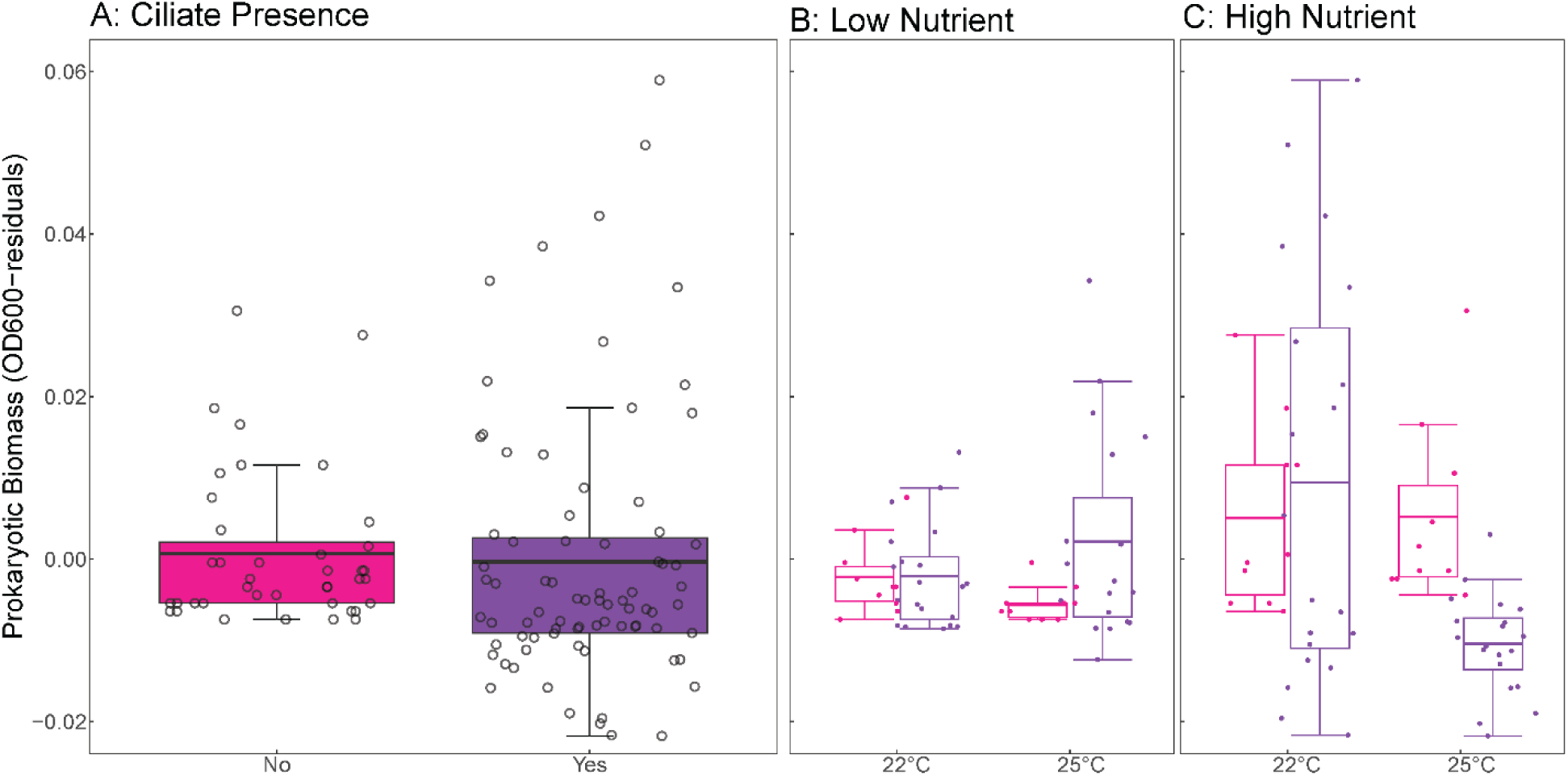
The independent and interactive effects of temperature, nutrients, and ciliate presence on prokaryotic biomass. (A) The presence of a ciliate community affects prokaryotic biomass. (B) Interaction between temperature, low nutrient levels, and ciliate presence. (C) Interaction between temperature, high nutrient levels, and ciliate presence. Each boxplot displays the mean (horizontal line), the 25th and 75th percentiles (box edges), and whiskers extending to the largest value within 1.5× the interquartile range (IQR). Open circles represent individual raw data points. Pink indicates treatments without ciliates; purple indicates treatments with ciliates.

The ciliate community did not influence total community respiration (effect = 9.601 × 10^-5^ ± 1.013 × 10^-4^ SE, t-value = 0.948, df = 112, p-value = 0.345, Fig 2). However, a weak two-way interaction between temperature and nutrients led to decreased respiration under high nutrient conditions at 25°C relative to other treatments (effect = 0.000331 ± 0.000165 SE, t-value = 2.004, df = 112, p-value = 0.048, Fig 2B, 2C), and no significant effect of ciliates presence (Fig 2B, 2C).

**Figure 2.**
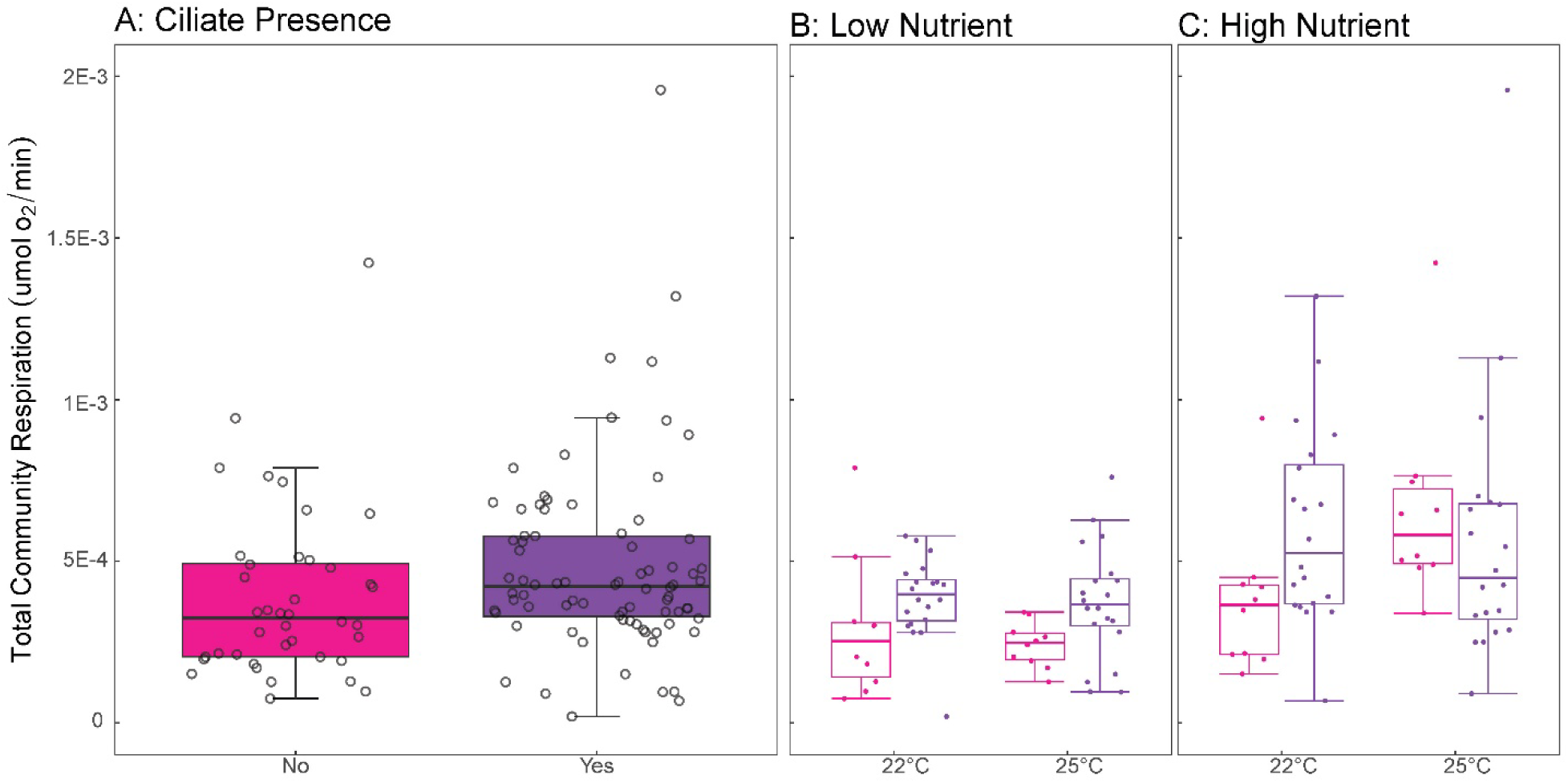
The independent and interactive effects of temperature, nutrients, and ciliate presence on total community respiration. (A) the presence of ciliate community effects on total respiration, (B) interaction between temperature, low nutrient level, and ciliate presence on total respiration, and (C) interaction between temperature, high nutrient level, and ciliate presence on total respiration. Lines within the boxplots represent median, 25th, and 75th percentile values, while whiskers are defined by the largest value not greater than 1.5× the interquartile range (IQR). Individual open circles represent raw data points. Pink colors represent treatments with no ciliates, while purple represents those that have the addition of ciliates.

### Direct effects of temperature, nutrients, and ciliate presence on the prokaryotic prey community diversity and composition

We quantified shifts in the Shannon diversity and composition of the prokaryotic community across treatments through 16S rRNA amplicon sequencing.

Proteobacteria were the most abundant group, followed by Firmicutes and Cyanobacteria (Figure 3A). Shannon diversity responded to a two-way interaction between temperature and ciliates (ANOVA, F = 7.3073, p-value = 0.007), as well as a three-way interaction between temperature, nutrients, and ciliates (ANOVA, F = 13.449, p-value = 3.76 × 10^-4^, Figure 3B) such that Shannon diversity increased in the presence of ciliates, but only under low nutrients and high temperatures (Figure 3B), leading to significant reduction in Firmicutes under those conditions relative to all other scenarios (Figure 3A).

**Figure 3.**
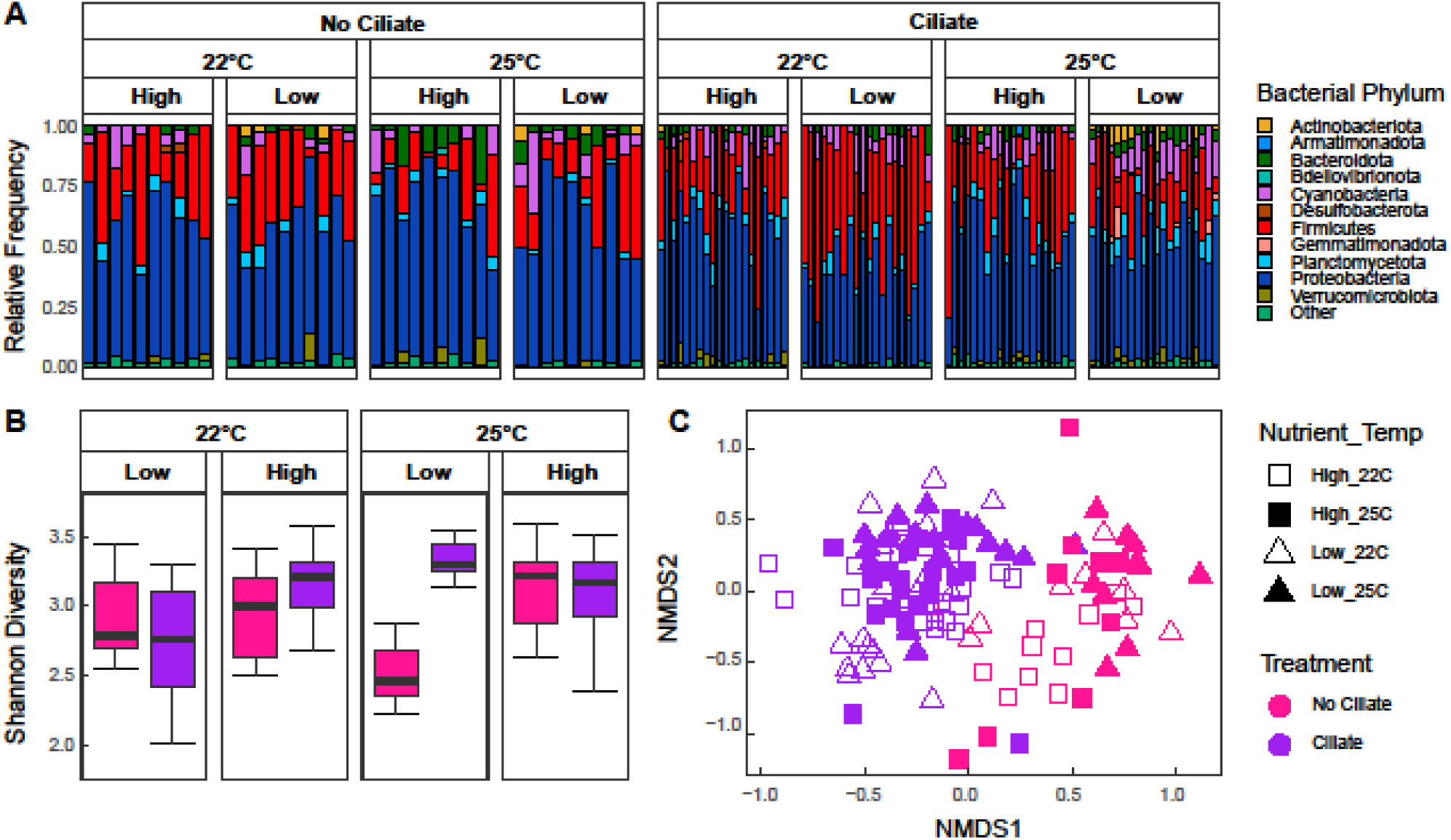
(A) Phyla relative abundances of prokaryotic (16S rRNA) prokaryotic communities across treatments. (B) Boxplot of alpha diversity of prokaryotic communities across temperature, nutrient level, and presence of ciliate community. Lines within the boxplots represent median, 25th, and 75th percentile values, while whiskers are defined by the largest value not greater than 1.5× the interquartile range (IQR). (C) NMDS ordination plot of Bray-Curtis dissimilarity of prokaryotic community composition (16S rRNA). Pink colors represent treatments with no ciliates, while purple represents those that have the addition of ciliates. Squares represent high nutrient conditions, while triangles represent low nutrient conditions. If the shape is filled in with color, it is experiencing warming (25), whereas those that are not filled in are at base conditions (22).

The composition of the prokaryotic prey community changed significantly due to two-way and three-way interactions between temperature, nutrients, and ciliate presence (Fig 3C). First, temperature increases species turnover under low nutrients but stabilizes it under high nutrient conditions (PERMANOVA, F = 2.491, p-value = 0.002, Fig 3C). Second, temperature amplified ciliate-driven effects on prokaryotic prey community composition (PERMANOVA, F = 2.056, p-value = 0.010, Fig 3C). Third, ciliates promoted higher species turnover under high nutrients than low ones (PERMANOVA, F = 1.970, p-value = 0.010, Fig 3C). Last, the ciliates strongly modulated the combined effects of temperature and nutrients on prokaryotic prey community composition (PERMANOVA, F = 2.336, p-value = 0.004, Figure 3C). For example, ciliates enhanced species turnover under high nutrient conditions at elevated temperatures, whereas their influence was weaker under low nutrient conditions regardless of temperature (Figure 3C).

### Direct effects of temperature and nutrients on the ciliate community

While temperature and nutrients directly influenced the prokaryotic community, ciliate presence also influenced their effect on prokaryotic biomass, diversity, and composition, but not total respiration. However, if the ciliates themselves respond to temperature and nutrients, the prokaryotic community could be responding indirectly to these ciliate shifts instead of simply showing different temperature and nutrient responses in their presence. To understand this, we first ask whether the ciliate communities respond to nutrients and temperatures by quantifying changes in ciliate density, diversity, and composition (see Methods).

Nutrients led to higher ciliate densities across all species, but this effect decreased with temperature (temperature: effect = 0.389 ± 0.192 SE, t-value = 2.025, p-value = 0.046; nutrient: effect = 0.865 ± 0.192 SE, t-value = 4.050, p-value = 2.4 × 10^-5^; two-way interaction effect = - 0.710 ± 0.271 SE, t-value = -2.613, df = 74, p-value = 0.010, Fig 4A).

**Figure 4.**
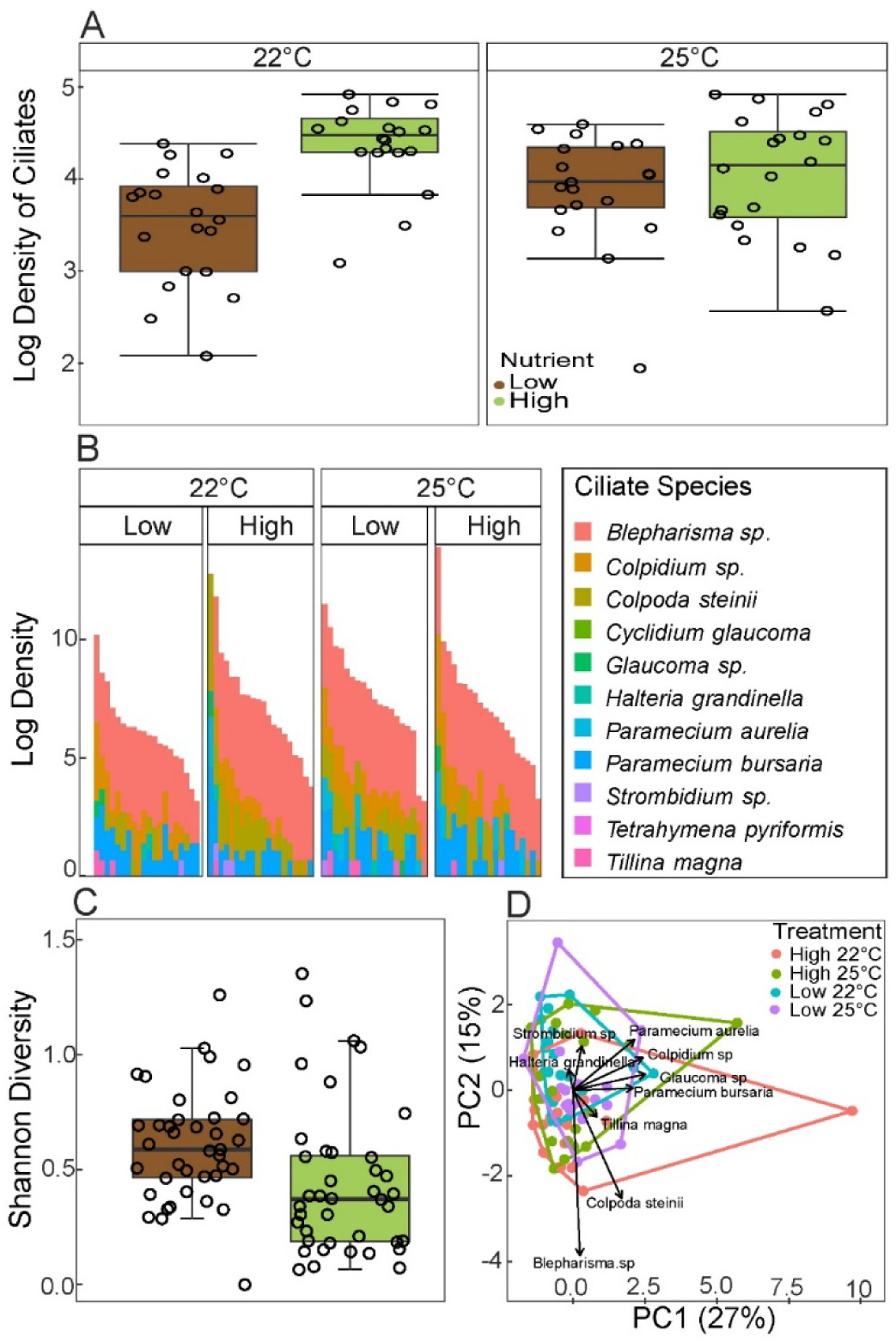
(A) Boxplot of ciliate density across temperature and nutrient level. Lines within the boxplots represent the median. (B) Final density data of ciliate communities for each sample across treatments. (C) Boxplot of alpha diversity of ciliate communities across temperature and nutrient level. Lines within the boxplots represent the median. (D) The ordination plot is based on a principal components analysis of ciliate community composition. Arrows show dominant communities’ variable loadings within the functional trait space, while individual points show individual samples and are colored by treatment. Individual open circles represent raw data points. Lines within the boxplots represent median, 25th, and 75th percentile values, while whiskers are defined by the largest value not greater than 1.5× the interquartile range (IQR).

*Blepharisma* sp. was the dominant ciliate species by density, followed by *Colpidium* sp. (Figure 4B); two species, *Tetrahymena pyriformis,* and *Cyclidium glaucoma,* were not present in any samples, most likely due to competition –or predation– by other ciliates (e.g., *Blepharisma* sp.). Ciliate Shannon diversity decreased with nutrients (effect = -0.229 ± 0.090 SE, t-value = - 2.542, df = 75, p-value = 0.013), and so did evenness (effect = -0.104 ± 0.041 SE, t-value = - 2.542, df = 75, p-value = 0.013), regardless of temperature (Fig 4C).

Ciliate communities shifted significantly in composition across temperature and nutrient treatments such that four distinct community configurations were possible––each corresponding to a principal component in PCA space: *Colpidium* sp.–*Paramecium bursaria–Paramecium aurelia–Glaucoma* sp. (PC1, Figure 4D), *Blepharisma* sp.–*Colpoda steinii* (PC2, Figure 4D), *Halteria grandinella*–*Strombidium* sp. (PC3, Figure 4D), *Tillina magna* (PC4, Figure 4D).

Composition was significantly affected by nutrients (PERMANOVA, F = 2.144, p-value = 0.021) and the interaction between nutrients and temperature (PERMANOVA, F = 2.022, p-value = 0.039).

### Direct and indirect effects of temperature and nutrients on the total microbial community structure and function

Having shown that temperature and nutrients directly affect the prokaryotic prey and ciliate predator community and that the presence of the ciliate community seems to mediate how the prey prokaryotic community responds to temperature and nutrients, we used a Structural Equation Modeling approach to understand the direct and indirect effects of temperature on the entirety of the microbial community. The best SEM model (model selection described in Appendix 1) converged after 11 iterations, including 25 parameters, and showed good alignment with our previous analyses while uncovering additional effects, both direct and indirect (Fig 5). All measures of goodness-of-fit suggested that the model correctly described the data (χ²(10) = 4.61, p-value = 0.990; CFI = 1.000; TLI = 1.194; RMSEA = 0.000, SRMR = 0.041).

**Figure 5.**
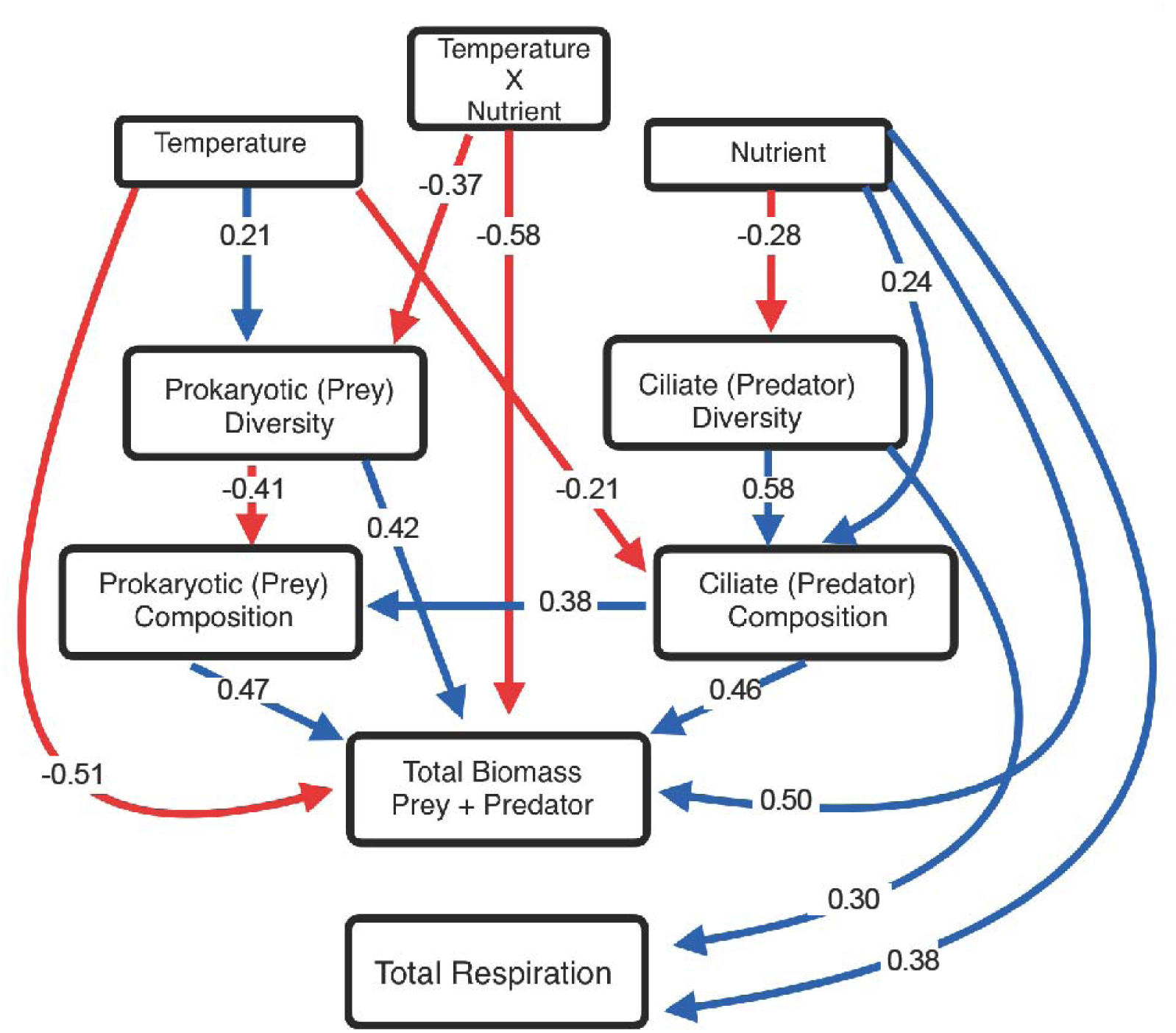
A structural equation model shows proposed relationships between latent variables, such as temperature and nutrients, and their interaction with the diversity, composition, and total biomass of microbes and their function. All numbers correspond to the standardized path coefficients. Solid arrows shown are significant direct effects (p < 0.05). Red arrows represent a negative relationship, while blue shows a positive.

In the best model, multiple regression paths were statistically significant, including a positive effect of temperature on the Shannon diversity of the prey prokaryotic community (β = 0.213, p = 0.036), a negative effect of nutrients on the Shannon Diversity of the predator ciliate community (β = -0.279, p = 0.009), and a negative effect of the interaction between nutrients and temperature on the prey prokaryotic diversity (β = -0.373, p < 0.001, Fig 5). Their diversity significantly influenced the composition of the predator ciliate and prey prokaryotic communities: prokaryotic diversity negatively influenced prokaryotic compositional change (β = -0.414, p < 0.001), while ciliate diversity positively influenced ciliate community compositional change (β = 0.58, p < 0.001). Nutrients led to ciliate community compositional change (β = 0.244, p = 0.024), while ciliate diversity declined with temperature (β = -0.210, p = 0.052). Last, the ciliate community composition drove prokaryotic composition (β = 0.375, p = 0.022), supporting all other analyses presented thus far (Fig 5).

Total microbial biomass –prokaryotic and ciliate– declined with temperature (β = -0.51, p = 0.001) and increased with nutrients (β = 0.50, p < 0.001). Prokaryotic diversity positively influenced total biomass (β = 0.34, p = 0.022), as did ciliate community composition (β = 0.464, p = 0.002) and prokaryotic community composition (β = 0.471, p = 0.003). A significant Temperature and Nutrient interaction (β = -0.58, p < 0.001) revealed that nutrient availability moderates the negative effect of temperature on total microbial biomass.

As opposed to biomass, total community respiration was influenced by fewer factors: nutrients led to an increase in total respiration (β = 0.382, p < 0.001), and so did ciliate diversity (β = 0.300, p = 0.004), suggesting that total respiration is more robust to shifts in the biotic and abiotic environment than composition or biomass. Last, total respiration was also negatively affected by nutrients indirectly through the diversity of the ciliates (β = -0.084, p = 0.052).

## Discussion

Understanding how temperature and nutrients influence microbial structure, biomass, and function is important in a warming world^35^. Our study shows how temperature and nutrients determine prokaryotic community biomass, composition, and respiration rates in the presence and absence of a ciliate community (Figures 1, 2, 3) and that the ciliate community also responds to these abiotic conditions (Figure 4). We then disentangle the direct and indirect effects of temperature and nutrients on the structure and function of the overall microbial community through their differential impacts on the diversity and composition of prokaryotes and ciliates, their biomass, and total respiration rates (Figure 5). Our results highlight the importance of ecological interactions in shaping prokaryotic community responses to environmental change.

We showed that elevated temperatures and nutrients reduced prokaryotic biomass (Figures 1B, 1C), aligning with observations in non-prokaryotic taxa across various ecosystems^45,73^. The presence of ciliates mediated these effects, likely due to selective predation on certain bacterial taxa^52^. While rising nutrient levels are expected to increase microbial biomass under high temperatures^27,74,75^, our findings do not support that trend, maybe owing to enhanced prokaryotic communities’ ability to maintain biomass despite nutrient fluctuations^76^, or compositional shifts, which were driven by interactive effects of temperature, nutrients, and ciliate presence (Figure 3C). Interestingly, ciliate presence drove species turnover in nutrient-rich environments (Figure 3C), which might be required for prokaryotic communities to maintain biomass production and function under shifting abiotic conditions, maybe resulting in a ciliate “rescue” effect of production and diversity under adverse conditions.

Total community respiration rates increased with temperature –consistent with theoretical expectations^77,5,7,46,78,38^– but only under high nutrient levels (Figures 2B and 2C), and showed no effect of ciliate presence (Figure 2A). This is striking, considering that predation by a single ciliate species was shown to decrease respiration under elevated temperatures^52^. Thus, the effects of a few species compared to entire ciliate communities can drastically affect their prokaryotic prey and their response to environmental change. While rising temperatures are expected to increase microbial respiration through increased heterotrophy^5,64,54^, our results suggest that may not always be true. This issue compounds with the fact that while ciliates showed density (Figure 4A, 4B), diversity (Figure 4C), compositional (Figure 4D), and biomass (Appendix 2) shifts with temperature and nutrients interactions consistent with laboratory^39^ and whole-ecosystem warming studies^72^, so neither nutrients nor temperature alone can fully explain the complex shifts in ciliate predator communities, possibly obscuring our ability to predict the fate of the entire microbial community under novel climates.

Our SEM approach indicated the direct and indirect effects of temperature and nutrients on the structure and function of the different components of the microbial community (Figure 5). For example, temperature directly reduces total microbial biomass. However, it also increases prokaryotic diversity, which negatively affects prokaryotic compositional variation from microcosm to microcosm, which increases total microbial biomass, resulting in an indirect effect that, while synergistic with its direct effect on total biomass, occurs through a completely different –indirect– mechanism. Interestingly, strong positive effects of ciliate community composition on prokaryotic compositional change highlight a possible role of top-down control in these microbial dynamics. In non-microbial food webs, the loss of top predators often mediates how the food web responds to abiotic shift conditions^79–81^, which our results suggest could also occur in our study system.

Accounting for both direct and indirect effects of abiotic conditions revealed that nutrients, but not temperature, influenced total microbial respiration directly and indirectly, through ciliate diversity and the effect of this diversity on respiration (Fig. 5). This highlights the importance of accounting for indirect effects when understanding joint biotic and abiotic effects on microbial function: indirect effects are likely to manifest as strong collinearity among explanatory model variables in a linear model framework, which often results in one variable’s effect masking the other’s (e.g., Fig. 2). Last, the conspicuous lack of explanatory variables for total community respiration –e.g., compared to total community biomass (Fig. 5)– suggests some level of robustness of respiration to shifts in community composition in response to environmental change^64,65,52^ and highlighting that functioning may be less vulnerable to environmental change than composition^82,83^.

*Caveats*. We did not track the dynamics of either community over time. Consequently, our results represent a snapshot in time, and some effects reported here may be transient, limiting our understanding of the underlying mechanisms and processes. As global temperatures rise, seasonality and variability are also changing^84,85^, but because we only manipulate mean temperature, our results do not inform how microbial communities may respond to shifts in temperature seasonality or variability and whether the effects of temperature variability are distinct from those of mean temperature, remains an open question. While the results presented are clear enough to suggest that these processes may also be at play in nature, the short-term nature of our experimental manipulations and tightly controlled experimental setup inevitably limit the possible scope of our inference.

*Conclusions*. Our results shed some light on how warming may affect carbon sequestration in peatlands. Indeed, peatlands are exceptionally susceptible to future climate change impacts^86–88^. Despite covering less than 3% of the Earth’s surface, these ecosystems store approximately 25%-30 % of the world’s soil carbon as recalcitrant peat^89^. However, peat moss growth in nutrient-poor peatlands depends on symbiotic interactions with prokaryotic associates ^90,91^. Shifts in prokaryotic community composition due to warming^92^ may impact the activity of moss symbionts and other important microbes, potentially leading to reductions in carbon sequestration in these peatlands through changes in moss growth^93^. Our results highlight that to fully understand the breadth and consequences of these changes, we need to consider the interactive effects of rising temperatures and increasing nutrient loads in these rapidly changing ecosystems. Moreover, our results suggest that essential but largely overlooked predatory interactions between these organisms and their predators may be important to understanding and predicting how climate change may affect the responses of these microbial communities in peatlands^13^. Still, total respiration levels might be influenced by a much narrower set of biotic and abiotic variables.

## Materials and Methods

### Prey Prokaryotic Community

We isolated a prokaryotic prey community from a peatland bog in the Marcell experimental forest (Minnesota, USA), next to the Spruce and Peatland Responses Under Changing Environment whole-ecosystem warming experiment^94^ from 5 cm core samples containing a top layer of living *Sphagnum* moss tissue and a lower layer of peat. We filtered out larger fungi and ciliates using 11μm pore size Whatman autoclaved filters, then removed smaller flagellates and fungal spores^48^ using sterile 1.6 μm pore size Whatman GF/A filters. While the community is not guaranteed to only have prokaryotes (e.g., eukaryotic nanoflagellates cannot be filtered out using this approach), it should be composed in its majority by Prokaryotes and Archaea^52^. From now on, we call it the *prokaryotic* community.

### Experimental Treatments

The prokaryotic community was homogenized and incubated in 120, 200mL acid-washed and autoclaved borosilicate jars filled with 150 mL of Carolina Biological Ciliate culture medium plus one autoclaved wheat seed as a carbon source^95^. Jars were assigned to nutrient to possible nutrient treatments –low nutrients (75 mL of media and 75 mL of distilled water, 60 jars) and high nutrients (150mL of media, 60 jars)– and one of two possible temperature treatments (60 jars at 22°C, representing ambient temperature, and 60 jars at 25°C, representing an increase in the average temperature of 3°C. There was no observable difference in pH between low and high nutrient treatments (pH measured using a Mettler Toledo SevenGo Duo SG68 pH/Ion/DO Meter). We added one autoclaved wheat seed (∼35 mg ea.) to high-nutrient treatments and half a seed (∼17.5 mg ea.) to the low-nutrient treatments^95,60^, both from the same batch to control for seed-associated microbes. All microcosms were incubated for three days in Percival AL-22 L2 growth chambers (Percival Scientific, Perry, Iowa) at 10% light intensity (1700 lux), 75% humidity, and a 16:8 hr. day-night cycle before further manipulation.

After three days, we added a synthetic ciliate community to 80 jars across all nutrient and temperature treatments, leaving 40 no-ciliate jars equally distributed among temperature and nutrient manipulations. This asymmetric design ensures the detection of even small effects in the jars with ciliates. We included the following ciliates: *Tillina magna, Tetrahymena pyriformis, Cyclidium glaucoma, Colpoda steinii, Strombidium* sp*., Blepharisma* sp.*, Glaucoma* sp.*, Colpidium* sp*., Halteria grandinella, Paramecium aurelia,* and *Paramecium bursaria* all of which naturally occur in *Sphagnum* peatlands^30^, and came from long-term (> five years) lab cultures^39^. Ciliates were introduced by pipetting well-mixed stock cultures at carrying capacity into experimental microcosms. Each species was introduced at a density of at least 17 individuals per jar (see Supplementary Table S6; Appendix 3 for initial densities). To control for possible effects of other prokaryotes in the ciliate cultures, we added filtered, homogenized ciliate growth medium to all "non-ciliate" jars, matching the amounts used in ciliate jars, as in previous studies^52,61,96^. The filtered media was inspected under the microscope to confirm the absence of ciliates before use, then confirmed again through microscopy and 18S sequencing post-experiment (no contamination was found). However, 18S sequencing revealed that one of the 80 ciliate jars had no ciliates by day 21 and was discarded for analysis.

Every jar was thus assigned to one of eight possible treatments –with 10 replicates for no ciliate jars, and 20 replicates for jars with ciliates: (1) 22°C, no ciliates, low nutrient; (2) 22°C, no ciliates, high nutrient; (3) 25°C, no ciliates, low nutrient; (4) 25°C, no ciliates, high nutrient; (5) 22°C, ciliates, low nutrient; (6) 22°C, ciliates, high nutrient; (7) 25°C, ciliates, low nutrient; and (8) 25°C, ciliates, high nutrient.

### Measurement and Analysis of Prokaryotic Biomass, Prey Community Diversity, and Total Community Respiration Rate

#### Prokaryotic Biomass

After 21 days, we quantified prokaryotic biomass, prey community diversity, and total community respiration rates. We measured the optical density at 600 nm wavelength (or OD600) as a proxy for the prokaryotic biomass^97^ using a BioTEK Epoch-2 spectrophotometer (Winooski, VT, USA). Higher OD600 indicates greater prokaryotic biomass^97^. Ciliates can scatter some light, but their contribution relative to that of the prokaryotes is much smaller^98,62^, due to their larger size and lower densities. Yet, OD600 could still reflect changes in ciliate biomass, even if it mostly reflects changes in prokaryotic biomass^98,62^. Our analyses show that ciliate biomass does a poor job at predicting OD600 (*R*^2^ = 0.14; Appendix 4), with a substantial spread away from the 1:1 line (SF1 Appendix 4), suggesting only a weak relationship between the two. To further control this minor influence, we used the residuals from a linear model fit to raw OD600 values as a function of ciliate biomass as a corrected estimate of prokaryotic biomass. These residuals (hereafter, prokaryotic biomass --OD600 residuals) were used in downstream models testing treatment effects. Based on these analyses and consistent with prior work^98,62^, we interpret OD600 as a reliable proxy for prokaryotic biomass rather than total microbial biomass.

#### Prey community diversity and composition

To estimate prokaryotic community diversity and composition, we transferred 1mL from all microcosms into 1.5mL Eppendorf tubes, pelleting and storing them at -80°C until DNA extraction. Genomic microbiome DNA was isolated with an Omega Mag-Bind Environmental kit. Amplification and preparation followed the Illumina 16S rRNA amplicon sequencing protocol with a custom mixture of 515F and 806R primers^99^ for archaea/bacteria and 18SV4F and 18SV4R primers^100^ targeting the 18S region. Prokaryotic sequences were processed with the QIIME 2 v 2021.2 platform^101^. Paired sequences were demultiplexed with the plugin demux and quality filtered (denoised, dereplicated, chimera filtered, and pair-end merged) and processed in Amplicon Sequence Variants (ASVs) with the DADA2 plugin^102^. Taxonomy was assigned using a pre-trained Naive Bayes classifier based on the Silva database (version 138) trimmed to the 515F/806R primer pair (16S rRNA). Sequence variant-based alpha (Shannon) and beta diversity (Bray-Curtis distance) were calculated with the *phyloseq* package^103^. Shannon diversity was calculated using the *diversity*() function in the R package "*vegan*" (v2.6-4;^104^). Beta diversity – which measures compositional change across jars– was calculated from a linear model test of the relationship between Bray-Curtis distance and jar.

#### Total Respiration rates

Total community respiration by day 21 was determined using optode-based real-time respirometry (OXY-4 SMA, PreSens, Regensburg, Germany)^95,105^ on the entire jar microcosm (150 ml) for 30 minutes and a collection rate of one measurement every three seconds, after a 30-minute acclimation period (n = 120) at their original experimental temperature and in the dark.

Respiration rates were estimated as the slope of the oxygen concentration over time (in μmol O_2_/min; Supplementary Figures S2, S3, and S4; Appendix 5)^106^.

### Analyses of Prokaryotic Biomass, Community Diversity and Composition, and Total Respiration Rates

Brute-force exploratory data analysis was performed by fitting all possible models with prokaryotic biomass (OD600) and respiration as response variables and all combinations of temperature, nutrients, ciliate presence, and their interactions as explanatory variables, then did multi-model inference to disentangle their joint effects –measured as the relative importance of each model term across all possible models– using R package MuMIn (v1.47.1)^107^. This analysis suggested that ciliate presence, temperature, and nutrient levels all interactively influenced prokaryotic biomass, while nutrients were the most important predictor of total microbial respiration, followed by the presence of ciliates and temperature (Supplementary Figures S6, S7; Appendix 6).

To quantify the magnitude and direction of these effects, we used separate linear models with ciliate presence, nutrient levels, temperature, and their two- and three-way interactions as explanatory variables, for either prokaryotic biomass (OD600), prokaryotic diversity (richness and Shannon diversity), or total respiration rates, as response variables. Additionally, we assessed how temperature, nutrients, and ciliate presence influenced the composition of the prokaryotic community using PerMANOVA on compositional data from Bray-Curtis. All response variables were log-transformed, and explanatory variables were treated as categorical. All analyses were done in R version 4.2.2^108^.

### Changes in the Ciliate Density, Biomass, Diversity, and Composition: Estimation and Analyses

Because the ciliate community could potentially respond to the imposed treatments, we tracked all ciliate species densities through fluid imaging of 1 mL subsamples of all microcosms (FlowCam, Fluid Imaging Technologies, Scarborough, ME, United States) at day 21. The FlowCam takes pictures of all censused cells and can use these pictures to estimate length, width, area, volume and other properties, thus providing estimates of density, and allowing us to calculate cell volume, mass, and hence, biomass. Ciliate biomass was calculated as the average body mass of each ciliate species times its density (i.e., ∑_Species_ N_*i*_ M_*i*_, where N is the density of species *i* and M is its average mass). The average mass was estimated by using the volume of each censused individual –i.e., volume of an ellipsoid, 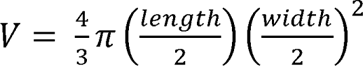, in µm³, and assuming that the two minor axes of each cell are the same. Individual cell volume was then converted into cubic centimeters (cm³) and then multiplied by the density of water (1 g/cm³) to individual cell mass estimates. These estimates were averaged within each species to obtain average mass, then multiplied by each species’ density and added across species to obtain ciliate biomass density in g/mL.

Classification of individual cell images into different species was done manually, for accuracy, but measurements were taken automatically by the FlowCam’s proprietary software. Shannon ciliate diversity was calculated using the *diversity*() function in the R package "*vegan*" (v2.6-4;^104^). Mean dissimilarity, a measure of community composition, was quantified using the Bray-Curtis distances among jars. Bray-Curtis dissimilarity was calculated using *vegdist* function in R package “*vegan”*. We then calculated mean dissimilarity row-wise: the resulting mean dissimilarity values were appended to the original dataset and treated as a measure of community average differences in community composition (or beta diversity), for each sample^109,110^.

We used linear models to evaluate how ciliate density and diversity changed with temperature, nutrients, or their interaction. We used Principal Components Analysis (PCA) on the ciliate community density data across all species, then tested for individual and interactive treatment effects on the first two principal components of such compositional data (PC1= 27%, PC2= 15%) using perMANOVA as implemented the adonis2() function in "*vegan*".

### Direct and Indirect Effects of Temperature and Nutrients

So far, we have separately addressed how prokaryotic communities respond to direct effects of temperature and nutrients with and without ciliates, and how ciliates may also respond to direct effects of these abiotic variables. However, it is unclear whether temperature and nutrients indirectly affect prokaryotes via ciliate responses or whether and how microbial community function (prokaryotes + ciliates) is driven by responses to temperature and nutrients by either prokaryotes, ciliates, or both.

We address these questions by fitting alternative Structural Equation Models (SEMs) in R package *lavaan* (version 0.6-18)^111^. The most complex model included the effects of temperature, nutrients, and their interaction, on biotic variables, Shannon diversity and Bray-Curtis composition for both prokaryotic and ciliate communities. The models also included the effects of these abiotic and biotic variables on (1) total microbial biomass—defined as a composite of prokaryotic biomass (OD600) and ciliate biomass—and (2) the joint effects of the biotic and abiotic variables and total microbial biomass, on total community respiration (Table S3, Appendix 1). We retained the best model by AIC and BIC (Appendix 1, Table S1-4).

## Data Availability

The sequence data have been deposited in GenBank SRA under accession PRJNA1095004; https://www.ncbi.nlm.nih.gov/bioproject/PRJNA1095004. All other data and code have been deposited in GitHub: https://github.com/kmd304/Predation-Mediates-TempXNut.git.

## Acknowledgments

We thank Ze-Yi Han, Christopher Kilner, and Daniel J. Wieczynski for their feedback. This work was supported by the US Department of Energy, Office of Science, Office of Biological and Environmental Research, Genomic Science Program Grant award number DE-SC0020362, NSF DEB award number 2224819, and a Simons Foundation Early Career Fellowship in Aquatic Microbial Ecology and Evolution number LS-ECIAMEE-00001588 to JPG. Additional support for prokaryotic and ciliate collection from SPRUCE, was supported by US Department of Energy (DOE), Grant/Award Number: DE-AC05–00OR22725 to DJW.

