## Appendix for "Predation by a ciliate community mediates temperature and nutrient effects on a peatland prey prokaryotic community"

**INDEX:**

**Appendix 1: SEM**……………………………………………………………………………...…2

**Appendix 2: Predator Community Diversity and Composition**…………………..…..………5

**Appendix 3: Initial densities of added ciliates**………………………………………………….7

**Appendix 4:** **Prokaryotic Biomass** …………………………………….…………………..……8

**Appendix 5: Estimation of Total Respiration rates**……………………………..……………10

**Appendix 6: Prokaryotic Biomass and Total Respiration Supplemental Results**.........……14

**Appendix 1: SEM**

Table S1: Table shows model fit indices for several Structural Equation Modeling (SEM) for model comparison.

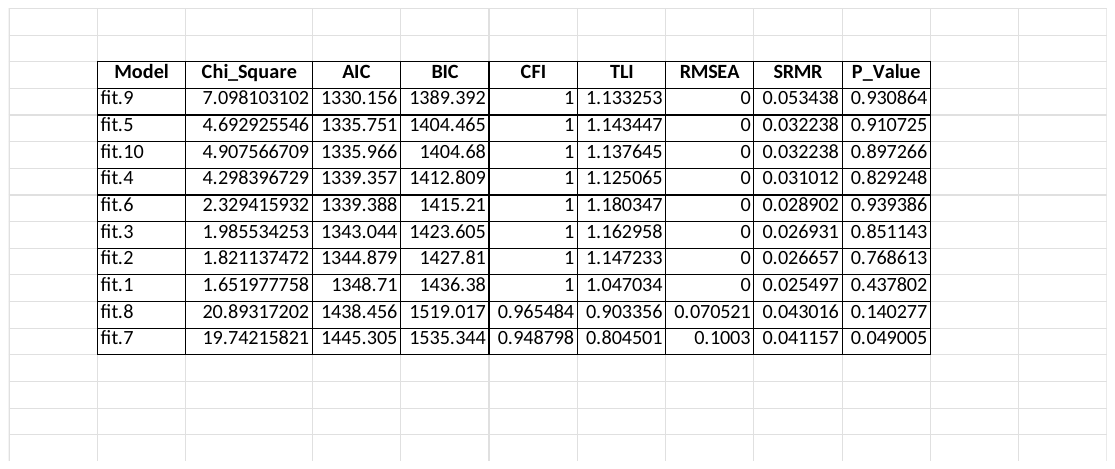

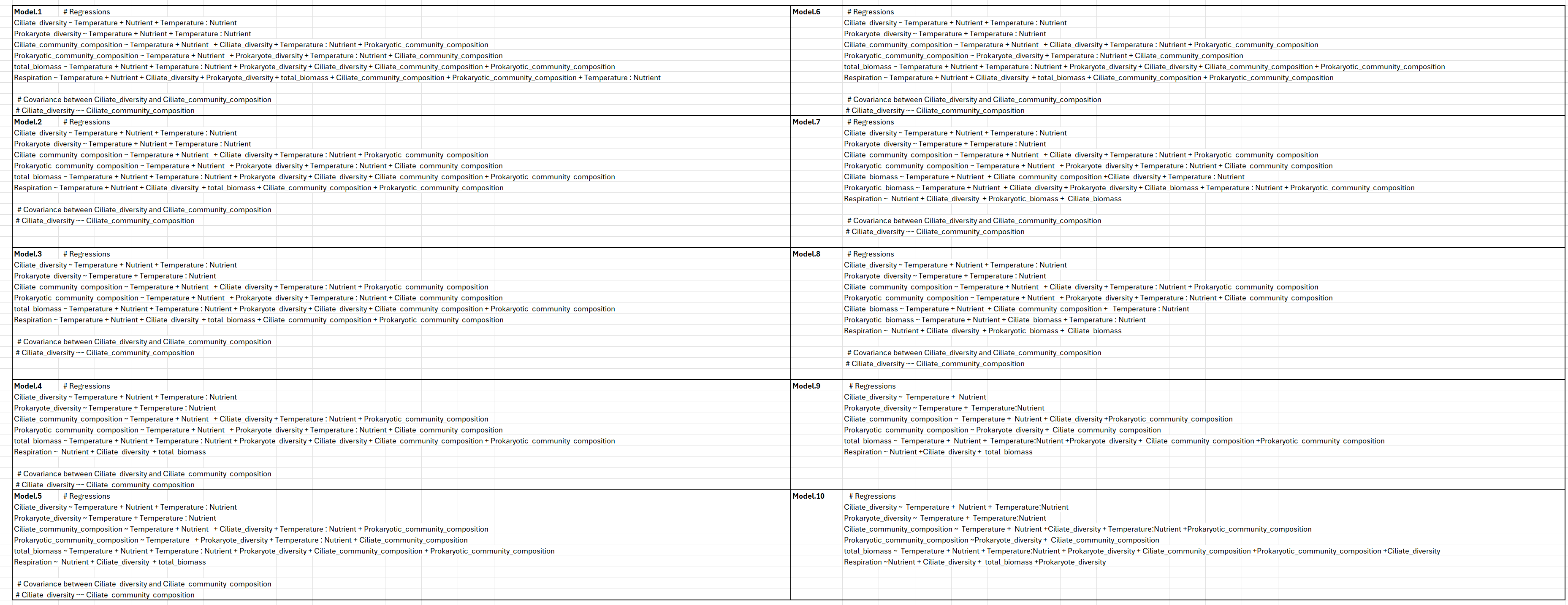
Table S2: Table shows all model structures for SEM.

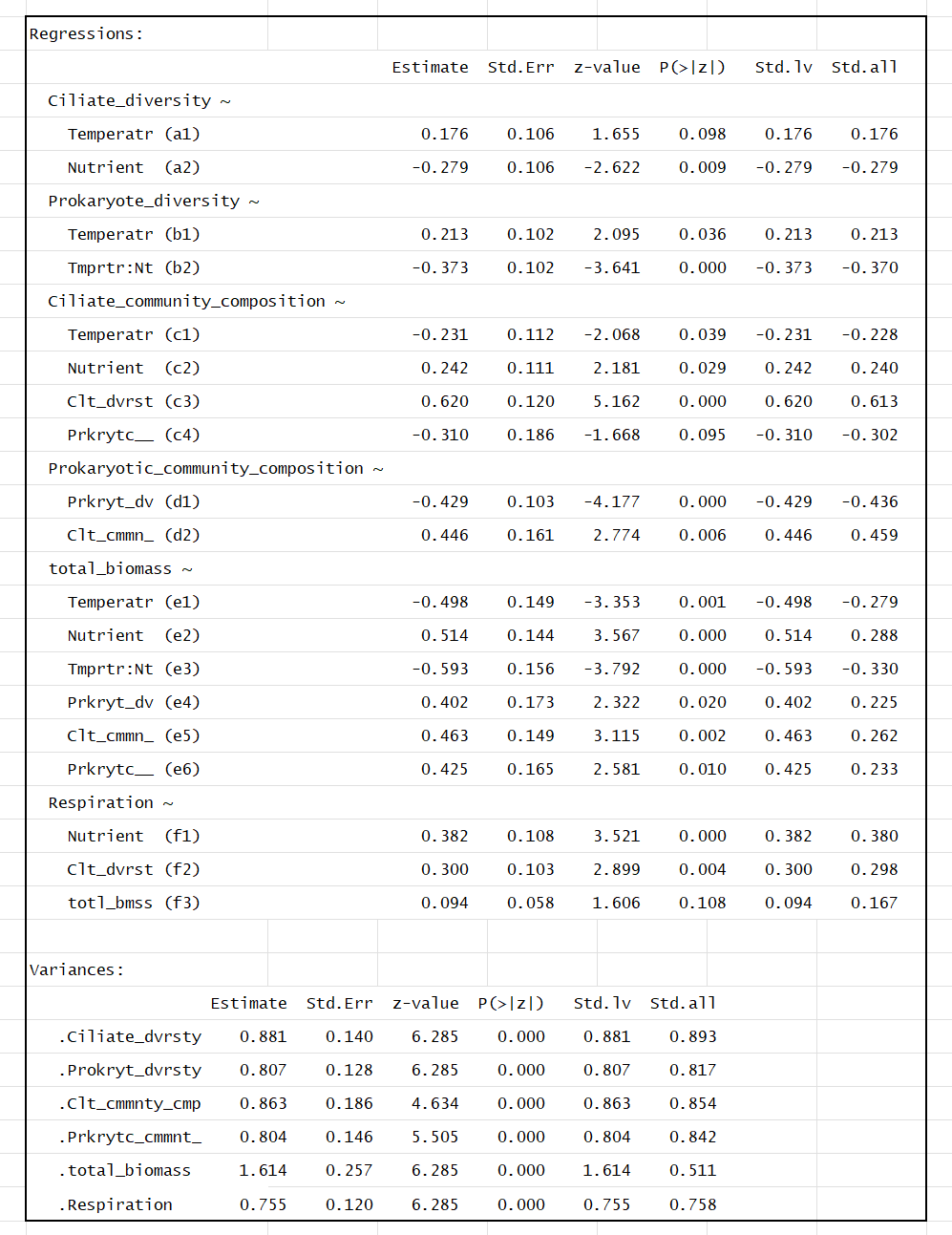
Table S3: Table shows the final model for SEM direct effects.

Table S4: Table shows the final model for SEM indirect effects.

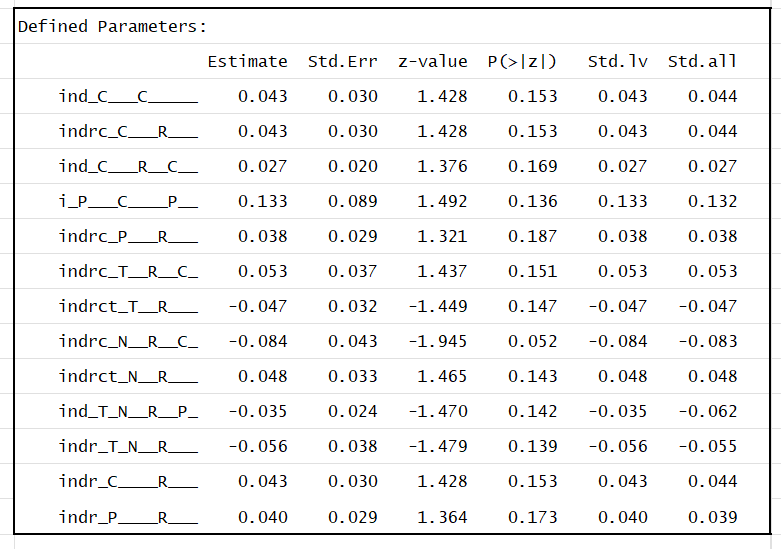

**Appendix 2: Predator Community Diversity and Composition**

Nutrients had a significant positive effect on ciliate densities (effect = 0.865 ± 0.1922 SE, t-value = 4.502, df = 74, p-value = 2.46 × 10^-4^). Temperature was associated with increased ciliate densities (effect = 0.3891 ± 0.1922 SE, t-value = 2.025, df = 74, p-value = 0.0465).

Effect of Temperature on Ciliate Biomass

The model shows that increasing temperature from 22°C to 25°C results in a significant decrease in total biomass by 1.063 × 10⁻⁵ g (p < 0.001). This suggests that microbial communities experience a negative effect when exposed to higher temperatures. One possible explanation is that elevated temperatures may increase metabolic demands, potentially leading to higher respiration rates and energy loss, which could reduce overall biomass accumulation. Additionally, certain microbial taxa may be less adapted to higher temperatures, leading to declines in their abundance and a subsequent reduction in total biomass.

Effect of Nutrient Availability on Ciliate Biomass

Reducing nutrient levels from Full to Half also leads to a significant reduction in biomass by 1.515 × 10⁻⁵ g (p < 0.001). This result aligns with ecological principles, as microbial growth is often limited by nutrient availability. When nutrients are scarce, microbial reproduction and metabolic activity slow down, leading to lower overall biomass accumulation. In resource-limited environments, competitive dynamics may also shift, with certain species outcompeting others, potentially leading to a decline in the overall microbial population.

Table S5: Table showing the results of the linear model of the effects of temperature and nutrients on ciliate biomass

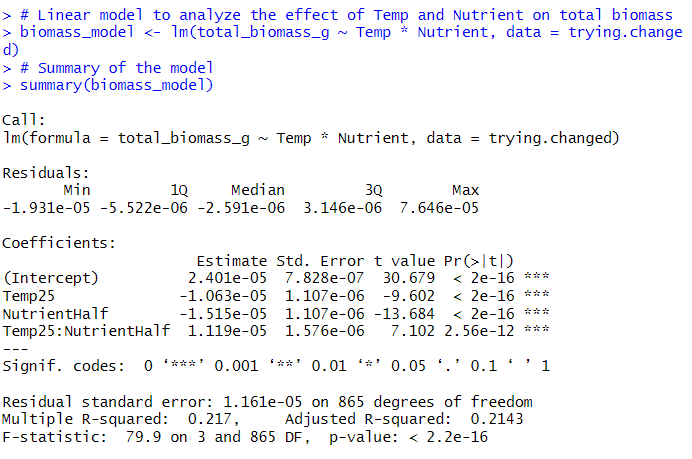

**Appendix 3: Initial densities of added ciliates**

**Table S6:** Table showing the density at which each ciliate species was introduced within the ciliate treatment jars.

| Species | Initial densities |
| --- | --- |
| Blepharisma sp. | 26in/mL |
| Colpidium sp. | 26in/mL |
| Colpoda steinii | 26in/mL |
| Cyclidium glaucoma | 17in/mL |
| Glaucoma sp. | 26in/mL |
| Halteria grandinella | 26in/mL |
| Paramecium aurelia | 17in/mL |
| Paramecium bursaria | 17in/mL |
| Strombidium sp. | 17in/mL |
| Tetrahymena pyriformis | 26in/mL |
| Tillina magna | 17in/mL |

**Appendix 4:** **Prokaryotic Biomass**

Z-score Transformation

To standardize the data for comparison, Z-scores were calculated for the variables OD600 and biomass.prot1. Z-score standardization transforms the data to have a mean of 0 and a standard deviation of 1, allowing for direct comparison between variables with different scales. Cook’s distance was used to identify influential points in the dataset. Cook’s distance quantifies how much an observation influences the regression model’s predictions. A threshold of 4/n (where n is the number of observations) was used to detect potential outliers. Observations exceeding this threshold were considered outliers and removed. After removing the outlier(s), a linear model was fitted to examine the relationship between Spec1_z (response variable) and biomass.prot1_z (predictor variable).

Table S7: Table showing the results of the linear model of

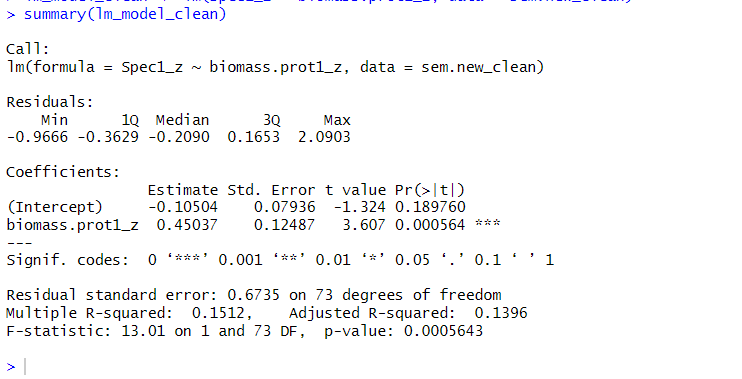

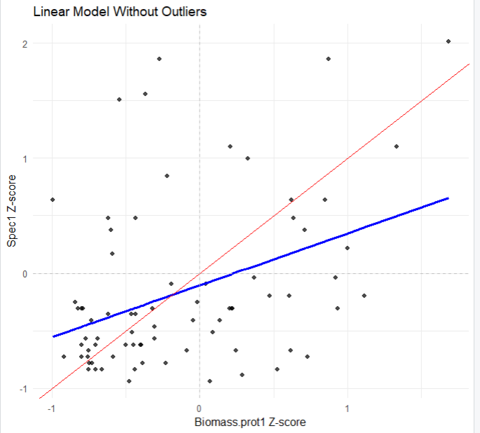

FS1. The scatterplot visualizes the relationship between Biomass.prot1 Z-score (x-axis) and OD600 (Spec1 Z-score) (y-axis) after removing outliers based on Cook’s distance. Each black dot represents an observation. The blue solid line represents the best-fit linear regression model, showing the predicted relationship between Biomass.prot1 Z-score and Spec1 Z-score. The red solid line represents a 1:1 relationship, meaning Biomass.prot1 Z-score was a perfect predictor of Spec1 Z-score; all data points would lie along this line.

Although there is a statistically significant correlation between protist biomass and OD600, with a highly significant p-value (p < 0.0005), the predictive power of this relationship is quite weak. The R² value of 0.16 indicates that only 16% of the variation in OD600 can be explained by protist biomass, meaning that the majority of the variation is driven by other factors not accounted for in this model. This is further illustrated by the scatterplot, where the data points form a large, dispersed cloud around the best-fit line rather than clustering tightly along it. The wide spread of values suggests that while protist biomass may have some influence on OD600, it does a poor job at accurately predicting it.

**Appendix 5: Estimation of total respiration rates**

Total respiration rates were measured as the slope (change over time) of the O2 concentration. Figs S2, S3, and S4 show the raw data for all jars and the linear regressions from which the slope (respiration rate in micro mol O2.L-1.min-1) was extracted.

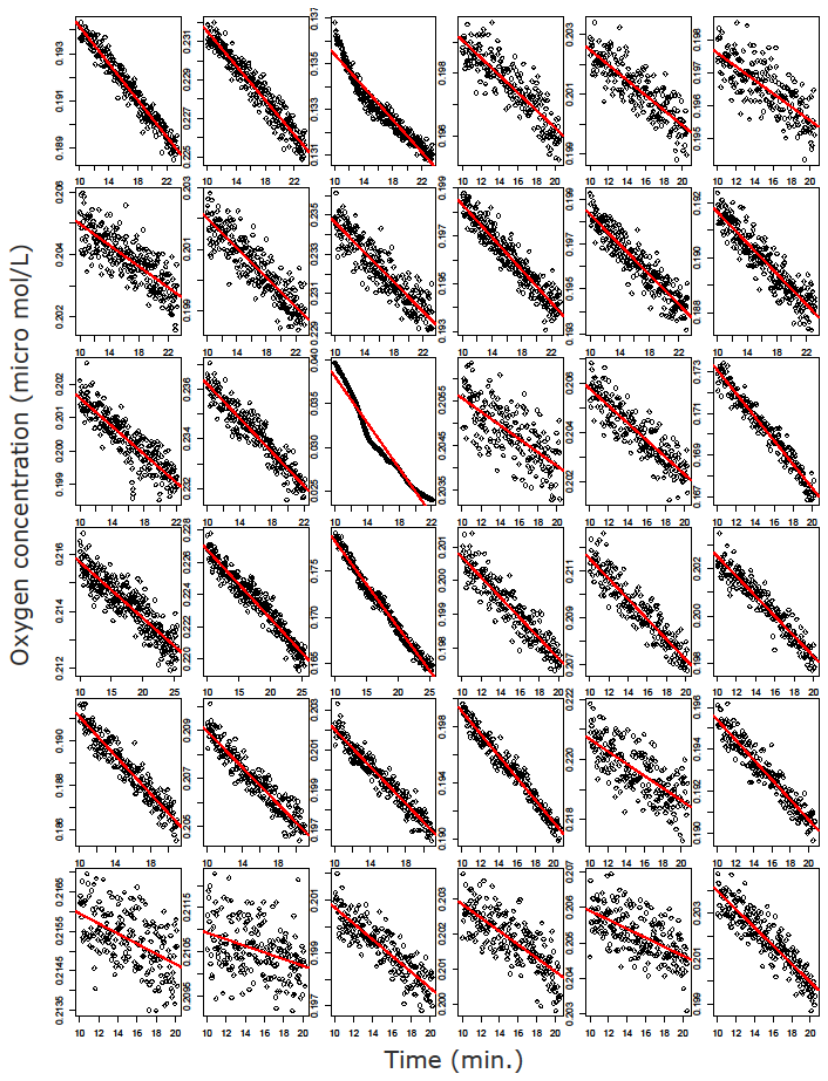

Fig S2: Raw data (grey) of all respirometry runs for day one. Red line represents the regression from which respiration rate (O2 consumed per unit time per L) was extracted.

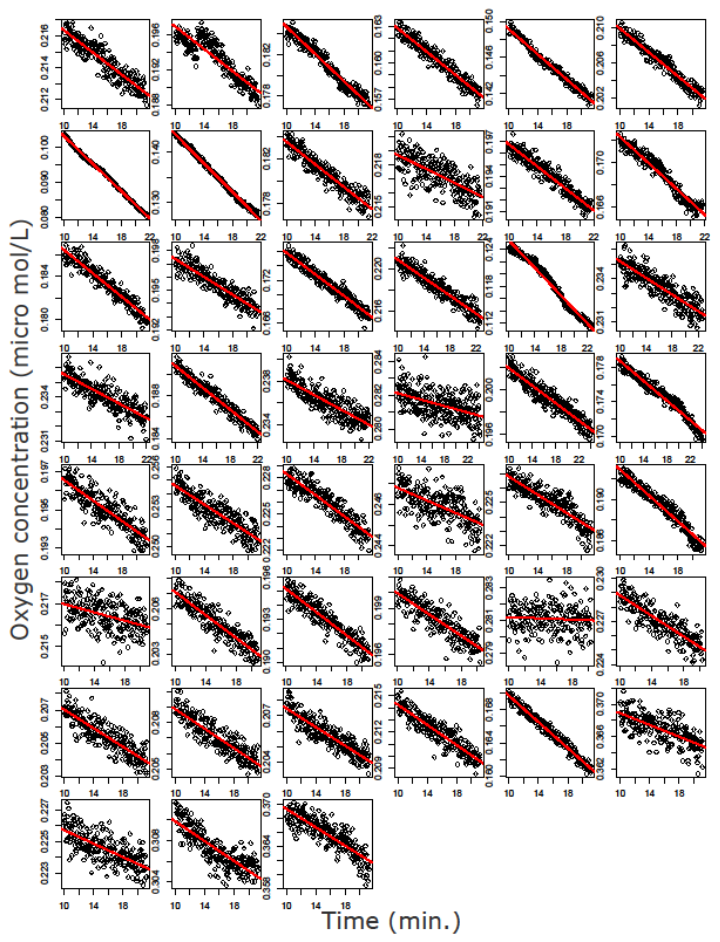

Fig S3: Raw data (grey) of all respirometry runs for day two. Red line represents the regression from which respiration rate (O2 consumed per unit time per L) was extracted.

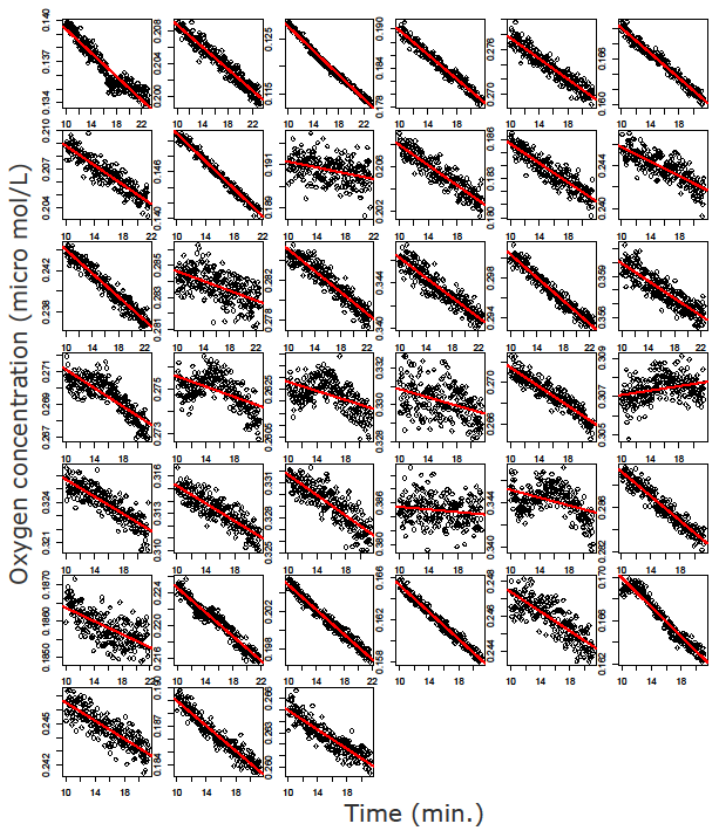

Fig S4: Raw data (grey) of all respirometry runs for day three. Red line represents the regression from which respiration rate (O2 consumed per unit time per L) was extracted.

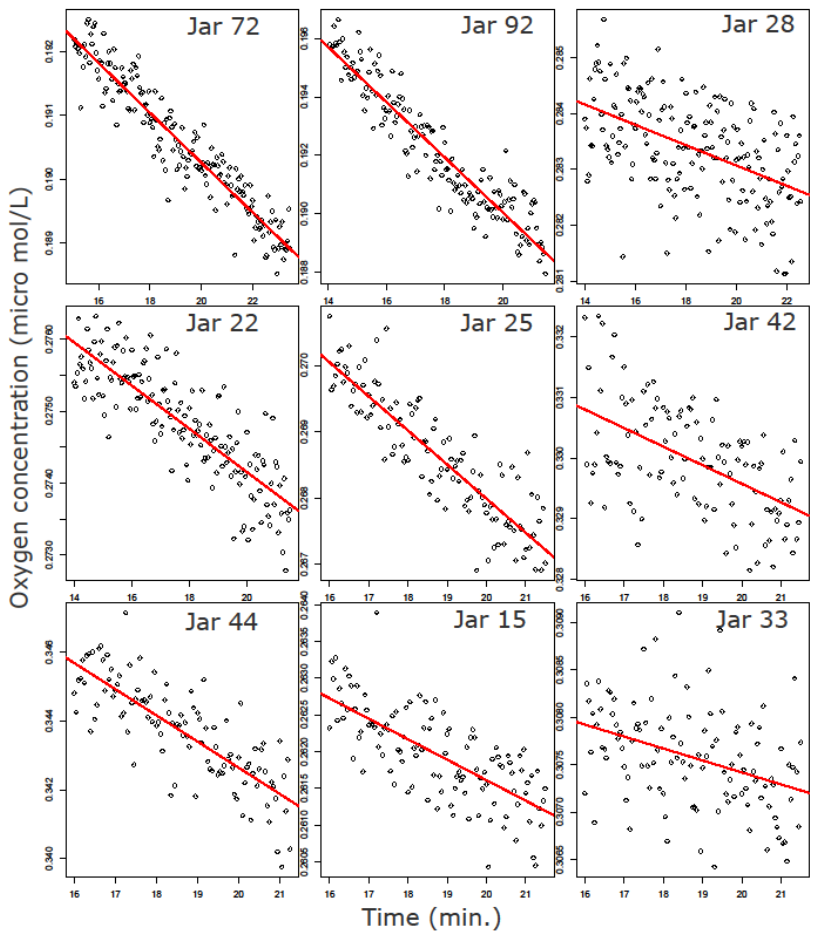

Fig S5: Curated data (grey) of nine respirometry runs. Red line represents the regression from which respiration rate (O2 consumed per unit time per L) was extracted.

**Appendix 6: Prokaryotic** **Biomass and Total Respiration Supplemental Results**

Prokaryotic Biomass (as OD600)

To understand the joint effects of Temperature, Nutrients, and Ciliates on prokaryotic biomass (OD600), we did exploratory data analysis using the R package “MuMIn”. In doing so, we fitted all possible models containing OD600 as the response variable and all combinations of Temperature, Nutrient, and Ciliate additions (No ciliates or ciliates) and their interactions as explanatory variables. This preliminary data exploration suggested that ciliate presence, temperature, and nutrient levels and all their interactions were important predictors of OD600, while nutrient levels were the most important for total respiration, followed by the presence of ciliates and temperature.

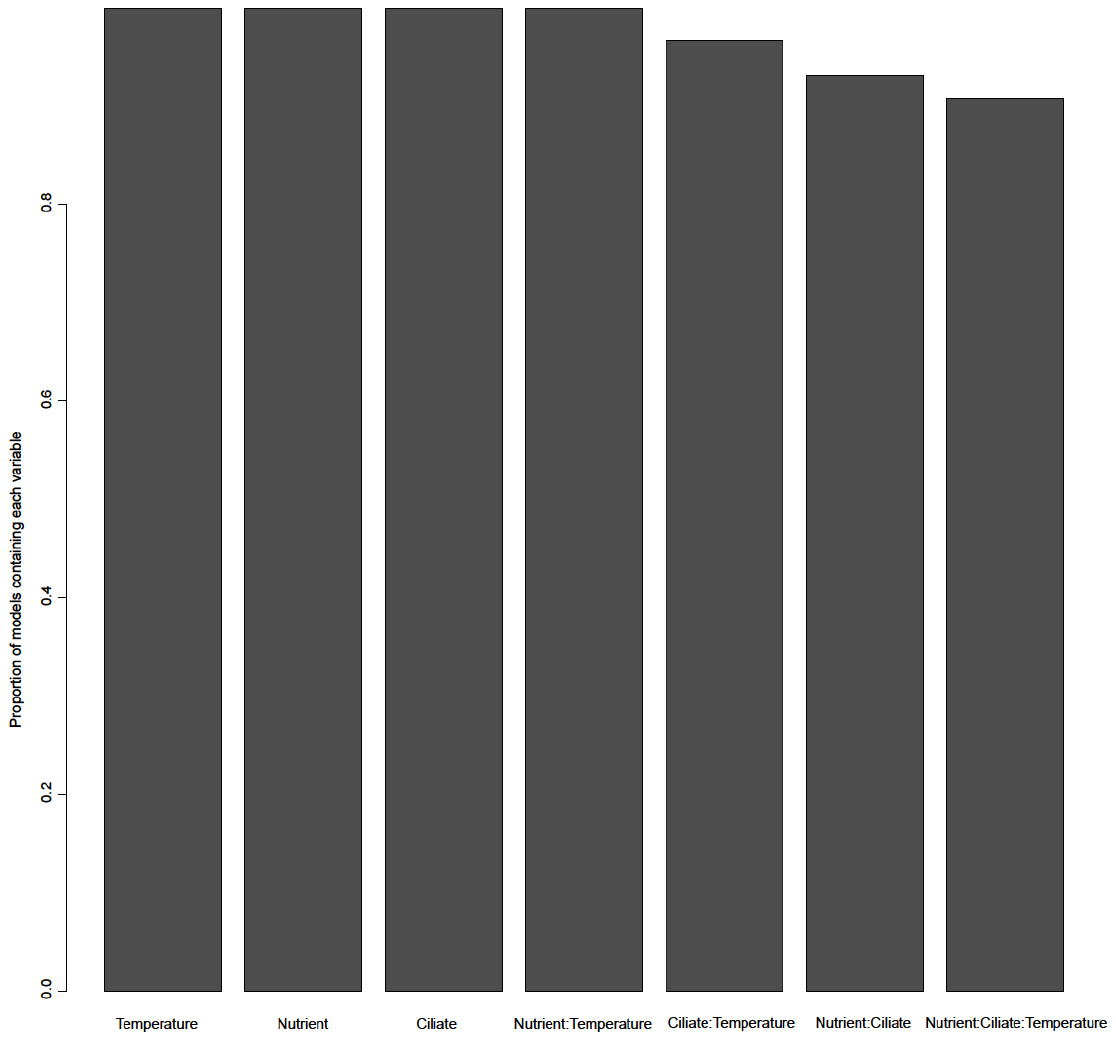

Fig S6 MuMIn ranks variables by how often they show up in models with good AICc support. Variables that show up in models with large AICc weight are considered to be more important than those that don’t. This figure shows how all variables are equally important.

Table S8: Table of the best models containing the variables temperature, nutrient level, ciliates, and all possible interactions ranked by AICc (output from “MuMIn” package). It can be seen that the best model accounts for Nutrients, Temperature, and Ciliates (Prt) and the interaction between nutrients and temperature. However, this model cannot be distinguished using AICc from one that only accounts for all variables and their interactions. This suggests that all these factors play an important role in determining OD600 (community biomass).

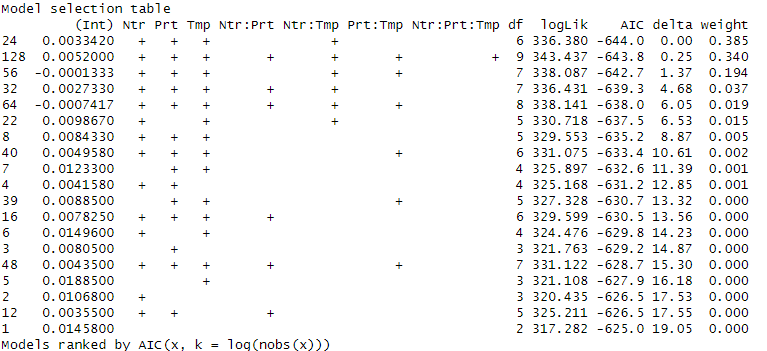

Table S9: Table showing the results of the linear model of interactive effects of temperature, nutrients, and ciliate presence on OD600 (prokaryotic community biomass).

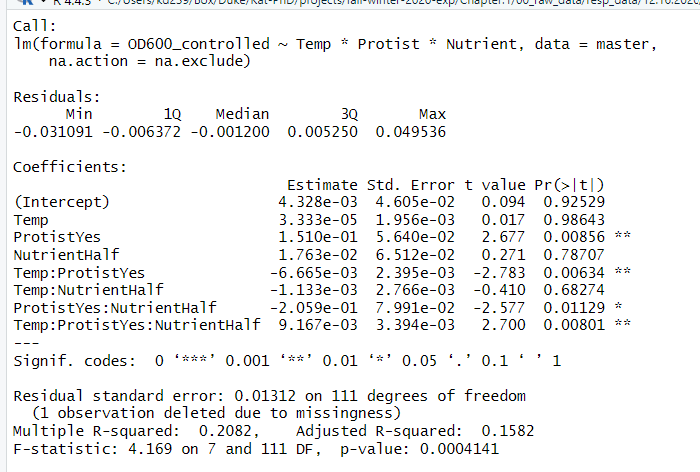

Table S10: Table showing the results of the linear model of independent effects of temperature on OD600 (prokaryotic community biomass).

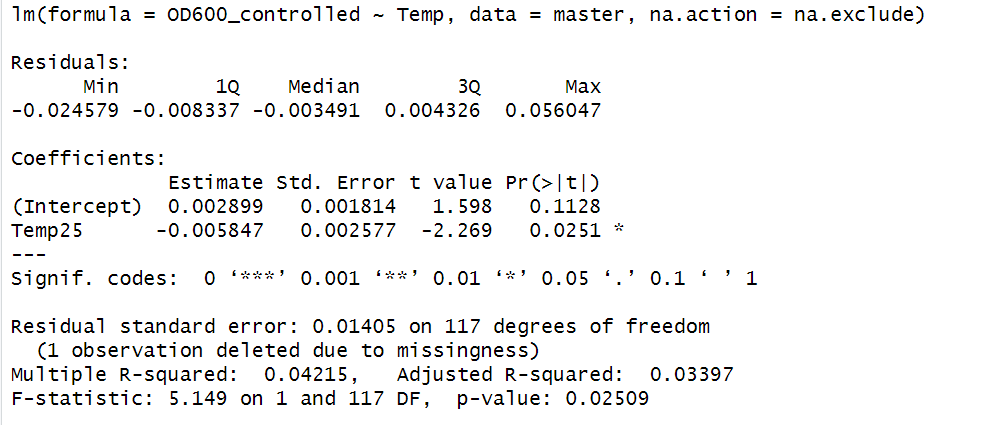

Table S11: Table showing the results of the linear model of independent effects of nutrients on OD600 (prokaryotic community biomass).

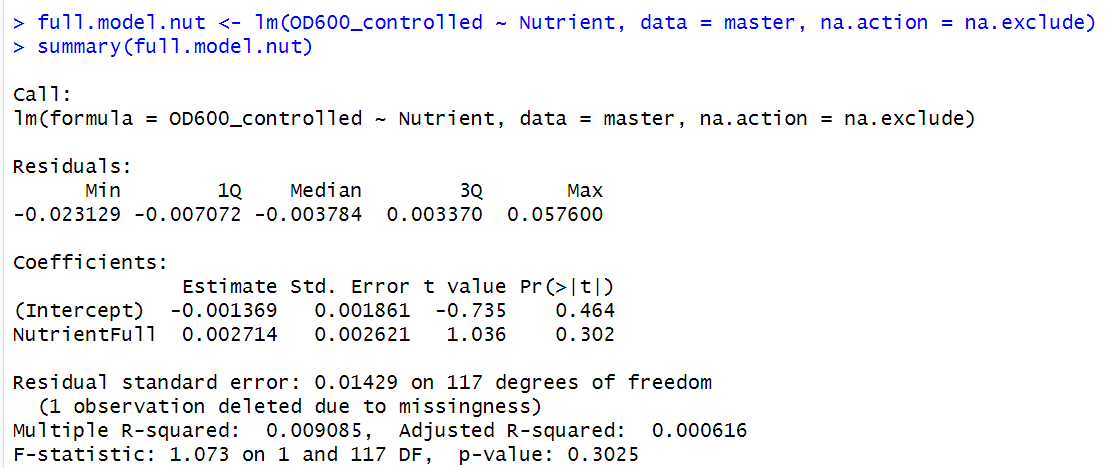

Table S12: Table showing the results of the linear model of independent effects of ciliate presence on OD600 (prokaryotic community biomass).

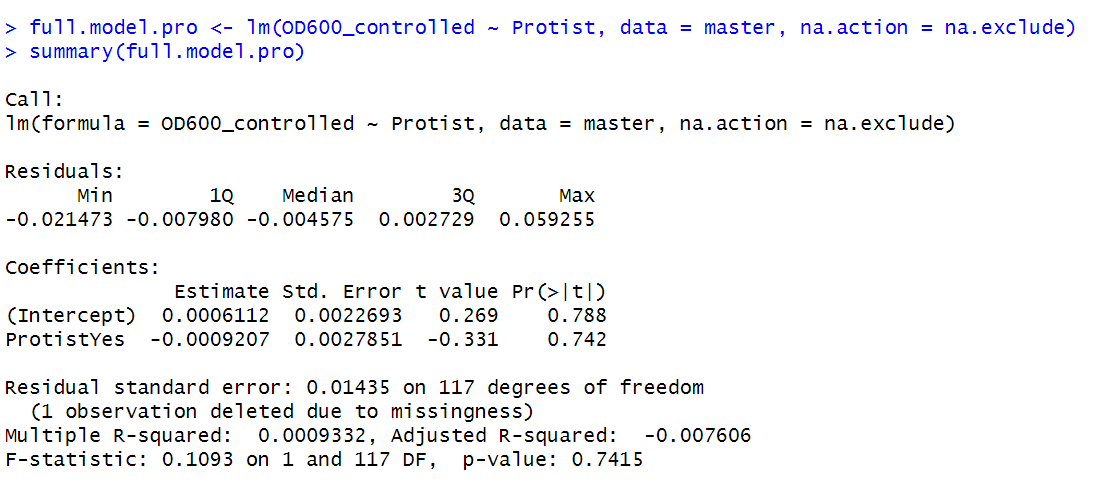

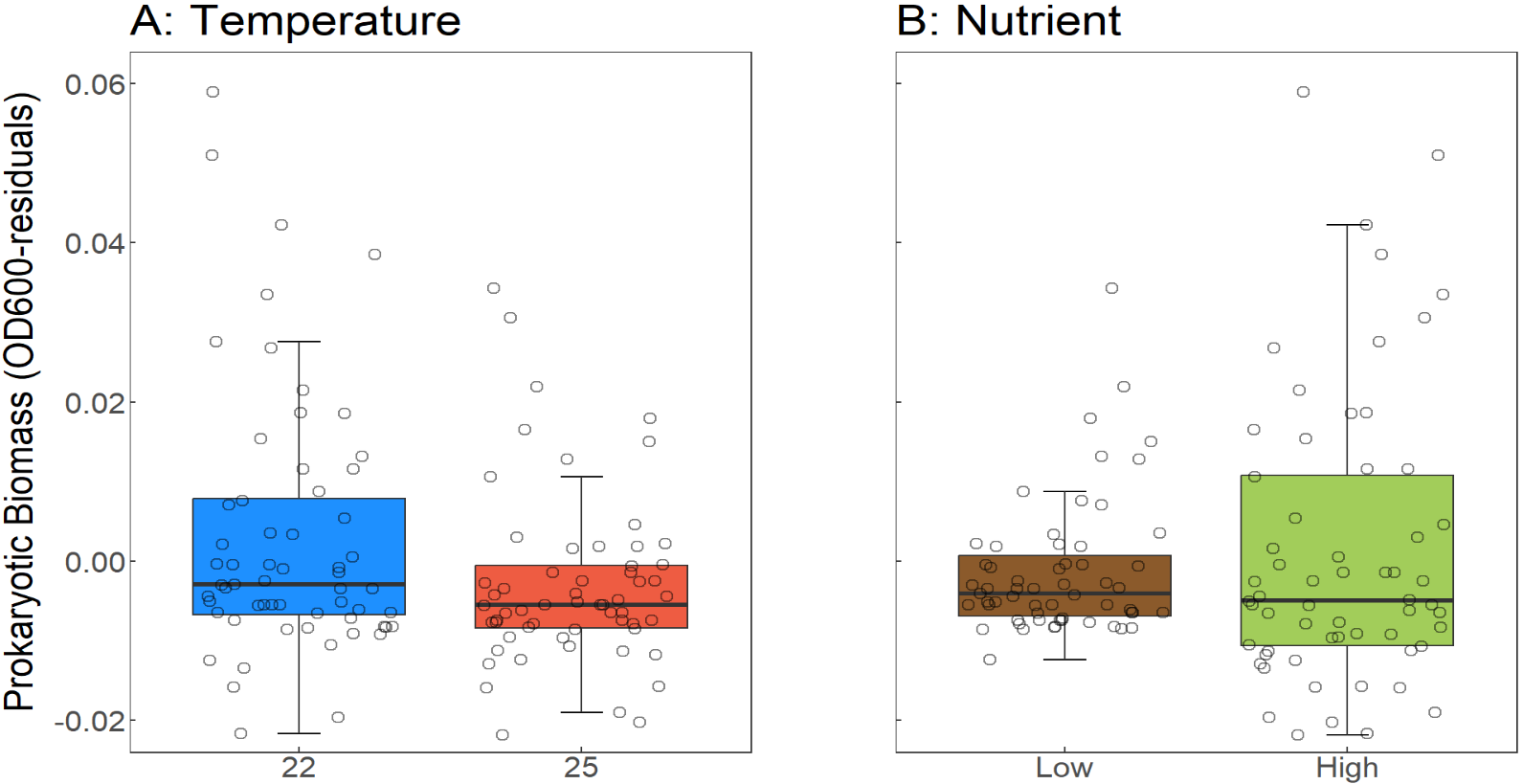

Fig S7. The independent and interactive effects of temperature, nutrients, and ciliate presence on prokaryotic biomass. (A) Boxplots of the temperature, (B) the nutrient level. Lines within the boxplots represent median, 25th, and 75th percentile values, while whiskers are defined by the largest value not greater than 1.5× the interquartile range (IQR). Individual open circles represent raw data points.

Respiration rates

Total respiration rates, such as prokaryotic Biomass (OD600), were analyzed. We, therefore, tested how all imposed treatments (Temperature, Nutrients, and Ciliates) influenced total respiration rates.

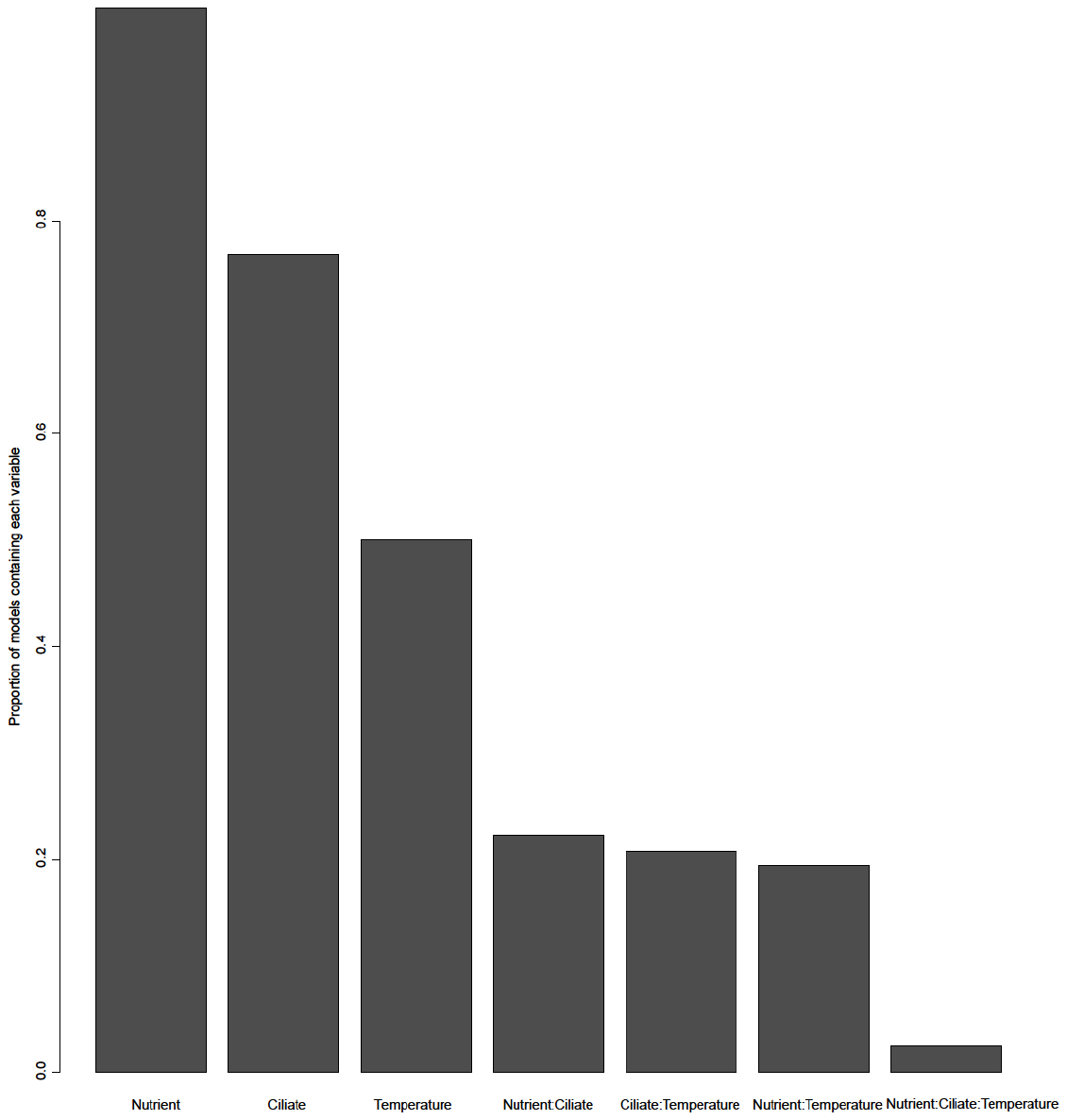

Fig S8: Clearly, the most important effects are those of nutrients, ciliates, and temperature, followed by all two-way interactions and then three-way interactions.

Table S13: Table of the best models containing the variables temperature, nutrient level, ciliates, and all possible interactions ranked by AICc (output from “MuMIn” package). It can be seen that the best model accounts for Nutrient level.

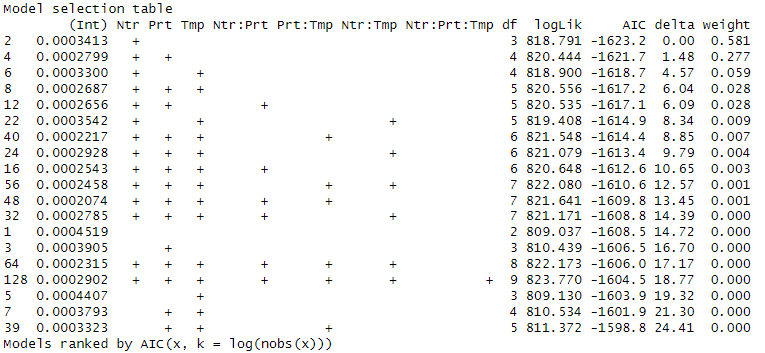

Table S14: Table showing the results of the linear model of interactive effects of temperature, nutrients, and ciliate presence on total respiration.

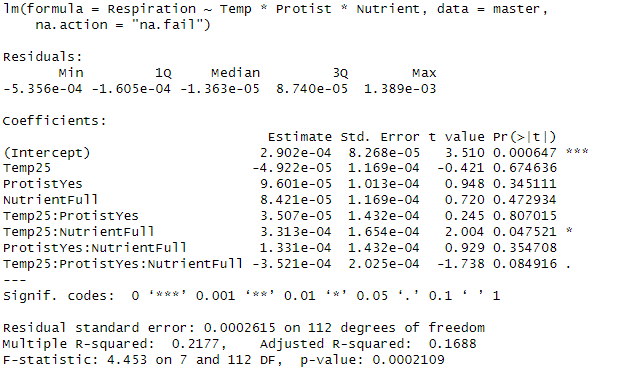

Table S15: Table showing the results of the linear model of independent effects of temperature on total respiration.

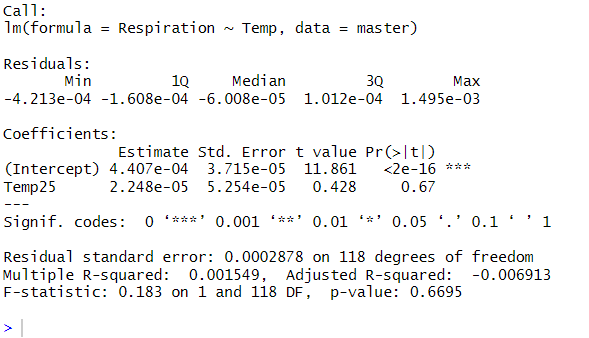

Table S16: Table showing the results of the linear model of independent effects of nutrients on total respiration.

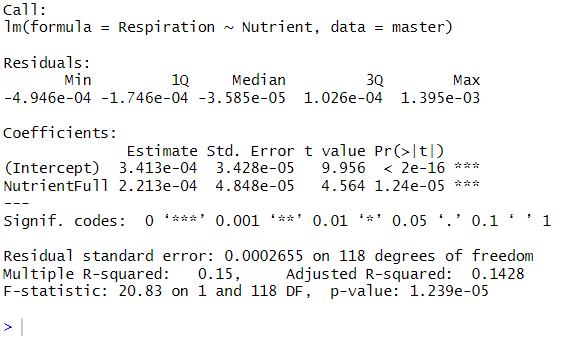

Table S17: Table showing the results of the linear model of independent effects of ciliate presence on total respiration.

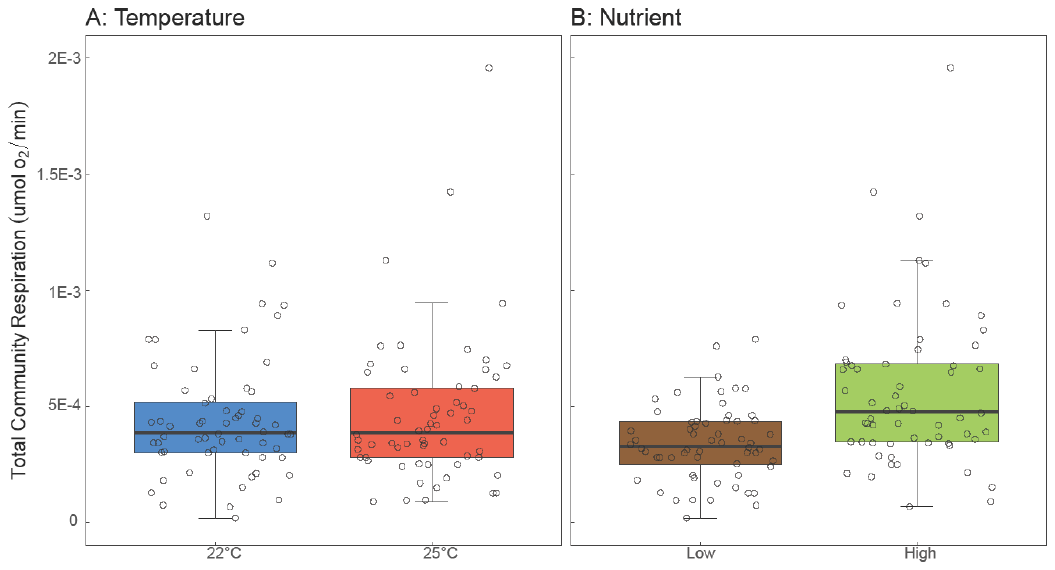

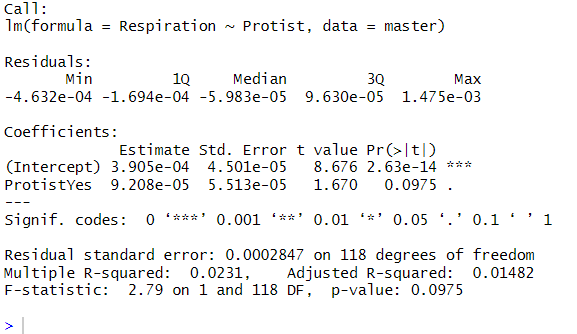

Fig S9. The independent and interactive effects of temperature, nutrients, and ciliate presence on total community respiration. (A) Boxplots of the temperature, (B) the nutrient level. Lines within the boxplots represent median, 25th, and 75th percentile values, while whiskers are defined by the largest value not greater than 1.5× the interquartile range (IQR). Individual open circles represent raw data points. Independent effects of nutrient levels were also significant. Total respiration rates were significantly higher under full nutrient conditions than low nutrient treatments. However, no significant direct effects of temperature or ciliate/predator presence on total respiration were detected.
